# Direct contact between iPSC-derived macrophages and hepatocytes drives reciprocal acquisition of Kupffer cell identity and hepatocyte maturation

**DOI:** 10.1101/2025.08.26.672295

**Authors:** Christopher Zhe Wei Lee, Farah Tasnim, Xiaozhong Huang, Raman Sethi, Yoohyun Song, Tatsuya Kozaki, Sebastiaan De Schepper, Nicholas Ang, Ivy Low, You Yi Hwang, Jinmiao Chen, Hanry Yu, Florent Ginhoux

**Affiliations:** Innovations in Food & Chemical Safety Programme (IFCS), A*STAR; Singapore Immunology Network, 8A Biomedical Grove, Singapore 138648, Singapore; School of Biological Sciences, Nanyang Technological University, Singapore 637551, Singapore; Institute of Bioengineering and Bioimaging, #05-01, 31 Biopolis Way, The Nanos, Singapore 138669, Singapore; Department of Physiology, Yong Loo Lin School of Medicine, National University of Singapore, MD9-04-07, 2 Medical Drive, Singapore 117593, Singapore; Bioinformatics Institute (BII), Agency for Science, Technology and Research (A*STAR), Singapore, Singapore; Institute of Molecular and Cell Biology (IMCB), Agency for Science, Technology and Research (A*STAR), Singapore, Singapore; NUS Graduate School – Integrative Sciences and Engineering Programme (ISEP), University Hall, Tan Chin Tuan Wing Level 5, #05-03, 21 Lower Kent Ridge Road, Singapore 119077, Singapore; Mechanobiology Institute, T-Labs, #05-01, 5A Engineering Drive 1, Singapore 117411, Singapore; CAMP IRG, Singapore-MIT Alliance for Research and Technology, 1 CREATE Way, Enterprise Wing, Level 4, Singapore, 138602, Singapore; Shanghai Institute of Immunology, Shanghai JiaoTong University School of Medicine, Shanghai, China; Translational Immunology Institute, SingHealth Duke-NUS Academic Medical Centre, Singapore, Singapore

**Keywords:** immune-mediated hepatotoxicity, *in vitro* liver models, co-culture, macrophages, Kupffer cells, induced pluripotent stem cells, iPSC, iMac

## Abstract

Hepatic macrophages play central roles in liver homeostasis, injury, and immune-mediated hepatotoxicity through dynamic crosstalk with hepatocytes. While monocyte-derived macrophages have been widely used in vitro, they do not fully recapitulate the biology of liver-resident Kupffer cells (KCs), which are embryonically derived and maintained locally. Recent advances suggest that induced pluripotent stem cell (iPSC)-derived macrophages (iMacs) more closely resemble embryonic macrophages and may therefore serve as a relevant platform to model KC biology.

Here, we developed a human iPSC-based co-culture system by combining iMacs with iPSC-derived hepatocytes (iHeps) derived from the same donor, enabling direct cell–cell interactions. We hypothesized that such interactions would both enhance hepatocyte maturation and promote KC-like differentiation of iMacs. Indeed, co-culture induced KC-like phenotypes in iMacs and improved functional maturation of iHeps, highlighting the importance of bidirectional cellular communication. Comparative analyses with iMacs cultured in hepatocyte-conditioned media revealed that direct contact provides additional signals beyond soluble factors in driving hepatic macrophage specialization. Functionally, this co-culture system demonstrated improved physiological relevance, particularly in modeling immune-mediated drug responses, as evidenced by enhanced cytokine production profiles upon exposure to a panel of test compounds.

Overall, this study establishes a novel human iPSC-derived platform that captures key aspects of hepatocyte–macrophage crosstalk, providing a more physiologically relevant model to investigate liver biology and assess immune-mediated drug toxicity.

## 1. Introduction

Hepatic macrophages play key roles in immune-mediated hepatotoxicity, liver injury and repair [1, 2]. They are impacted by autocrine and paracrine signals from hepatic cells, especially hepatocytes. They also release soluble stress signals in response to external stimuli from foreign particles, leading to macrophage activation and production of cell signalling and stress pathway modulators, including reactive oxygen species and cytokines [3, 4]. In return, hepatic macrophage activation also regulates parenchymal and non-parenchymal cell death and the metabolic activity of hepatocytes [5]. Such crosstalk between liver parenchymal cells (hepatocytes) and non-parenchymal modulators (hepatic macrophages) is critical in modelling liver injury, especially immune-mediated drug-induced responses [6].

Two major populations of hepatic macrophages have been reported: liver-resident Kupffer cells (KCs) and monocyte-derived macrophages (MoMLJs) that infiltrate the liver upon insult [3]. While MoMLJs have been well studied *in vivo* and applied in *in vitro* models [6], in more recent years it has been revealed that KCs are derived from embryonic macrophages that seed the liver early on during development [7][8], and are only minimally replaced by circulating monocytes under steady state conditions [9]. This also implies that human peripheral blood monocytes, the conventional source of primary human macrophages, might not be able to fully recapitulate KC biology. However, it has been demonstrated that murine iPSC-derived macrophages (iMacs) more closely resembled embryonic macrophages than monocyte-derived macrophages [10], suggesting that human iMacs might also serve as a suitable analogue for human embryonic macrophages. Based on these new insights, in our previous work, we differentiated iMacs into iPSC-derived KCs (iKCs) using primary hepatocyte conditioned media (PHCM) to drive iMacs towards iKC identity [11]. Although these iKCs partially resembled primary human KCs (PhKCs) based on the expression of KC specific markers and drug response, PHCM is not readily available and might exhibit differences depending on donor variability of primary hepatocytes. Most importantly, the direct cellular interactions between iMacs and hepatocytes and their potential to impart better reciprocal identity and functionality to each other has not been studied before.

We recently generated macrophage-sufficient brain organoids by coculturing brain organoids with iMacs generated from the same human iPSC line [12]. In such cocultures, iMac differentiated into cells with microglia-like phenotypes and functions. Most importantly, iMac modulated neuronal progenitor cell (NPC) differentiation, limiting NPC proliferation and promoting axonogenesis, profoundly remodelling organoid physiology. Thus, in this study, we hypothesized that iMacs could be directly cultured with iPSC-derived hepatocytes (iHeps) from the same IPSC source to both improve iHep development and maturation as well as impart KC-like identify to the iMacs. Upon development of such a model, we tested its application in detecting immune-mediated responses of seven paradigm compounds. Concurrently, we also cultured iMacs in PHCM for 7 days in a similar way to our previous study [11], comparing the transcriptomic effect of direct contact against soluble factors in the context of hepatic macrophage specialisation. Our results showed improved *in vivo* correlation based on the effects on cytokine production when iMac-derived KCs were used. Altogether, this study highlights the effect of direct crosstalk between hepatocytes and macrophages *in vitro* using iPSCs and provides a new human model to test drug-induced cytokine responses.

## 2. Materials and Methods

### 2.1 Maintenance of iPSCs

Human IMR90-iPSCs were obtained from WiCell Research Institute (Madison, WI). The cells were maintained under antibiotic-free conditions in mTESR media (StemCell Technologies, 85850) on Matrigel (Corning, 354234)-coated plates. Cells were passaged using ReLESR (StemCell Technologies, 05872) or Dispase (StemCell Technologies, 07913) as per manufacturer’s protocol whenever they reached 80% confluency.

### 2.2 Differentiation of iPSCs to iHeps

IMR90-iPSCs were differentiated into iPSC-hepatocytes (iHeps) as previously described [13]. Briefly, the iPSCs were cultured to 80% confluency, then made into single cells using Cell Dissociation Buffer (Gibco, 13151014) and seeded at 1.5 x 10^5 cells/cm^2^ on the desired well plate format (i.e. 1.5 x 10^6 cells for a 6 well plate) and placed under hypoxic conditions (5% O_2_ and 5% CO_2_). Growth factors and media used starting from the next day to day 20 are detailed in [11]. Media was changed every day and hypoxic conditions were used during days 5 to15 of differentiation. After 20 days of differentiation, iPSC-derived macrophages (iMacs) were either directly added to the iHeps as described in section 2.5 or iHeps were harvested for seeding into smaller well formats for drug testing. The harvesting protocol has been previously optimized by us and has been detailed in [14].

### 2.3 Differentiation of iPSCs to iMacs

IMR90-iPSCs were differentiated into iMacs as previously described [10]. Briefly, the iPSCs were cultured to 80% confluency, then passaged using ReLESR. As ReLESR releases the cells as small cell clumps, the passage conditions were optimised to result in roughly 1 x 10^5 cells in each well of a 6 well plate (Day-1). Starting from the next day (Day 0) to day 16, the cells were cultured in Stempro Media, consisting of Stempro-34 SFM (GIBCO, 10639-011), supplemented with 200 ug/mL Human Transferrin (Roche, 10-652-202-001), 1x L-Glut, 1x Pen/Strep, 0.5 mM Ascorbic Acid (Sigma, A4544) and 0.45mM MTG (Sigma, M6145). A full media change was done every other day for the next 16 days, supplemented with the following cytokines: Differentiation Day 0 (5 ng/mL BMP4, 50 ng/mL VEGF, and 2 uM CHIR99021), Differentiation Day 2 (5 ng/mL BMP4, 50 ng/mL VEGF, and 20 ng/mL FGF2), Differentiation Day 4 (15 ng/mL VEGF and 5 ng/mL FGF2), Differentiation Day 6 to 10 (10 ng/mL VEGF, 10 ng/mL FGF2, 50 ng/mL SCF, 30 ng/mL DKK-1 (RnD, 5439-DK), 10 ng/mL IL-6 (RnD, 206-IL), and 20 ng/mL IL-3), Differentiation Day 12 and 14 (10 ng/mL FGF2, 50 ng/mL SCF, 10 ng/mL IL-6, and 20 ng/mL IL-3). From day 16, the cells were cultured in 75% IMDM with Glutamax (GIBCO, 31980-030), 25% F12 (GIBCO, 11765-047), 1x N2 supplement (GIBCO, 17502-048), 1x B27 Supplement (GIBCO, 17504-001), 0.05% BSA (GE Healthcare, SH30574) and 1x Pen/Strep, supplemented with 50ng/ml of CSF-1 (RnD, 216-MC), with a full media change every 3 days. In addition, for the first 8 days of differentiation, the cells were kept in a hypoxic incubator (5% O_2_ and 5% CO_2_), before being transferred to a standard cell culture incubator for the next 18 days. After transfer to a standard cell culture incubator, floating cells were collected every media change and resuspended back into the culture to retain the hematopoietic progenitors. After 26 days, the floating cells were collected, and the purity of the iMacs was determined by flow cytometry (Supp Fig 1E).

**Figure 1.**
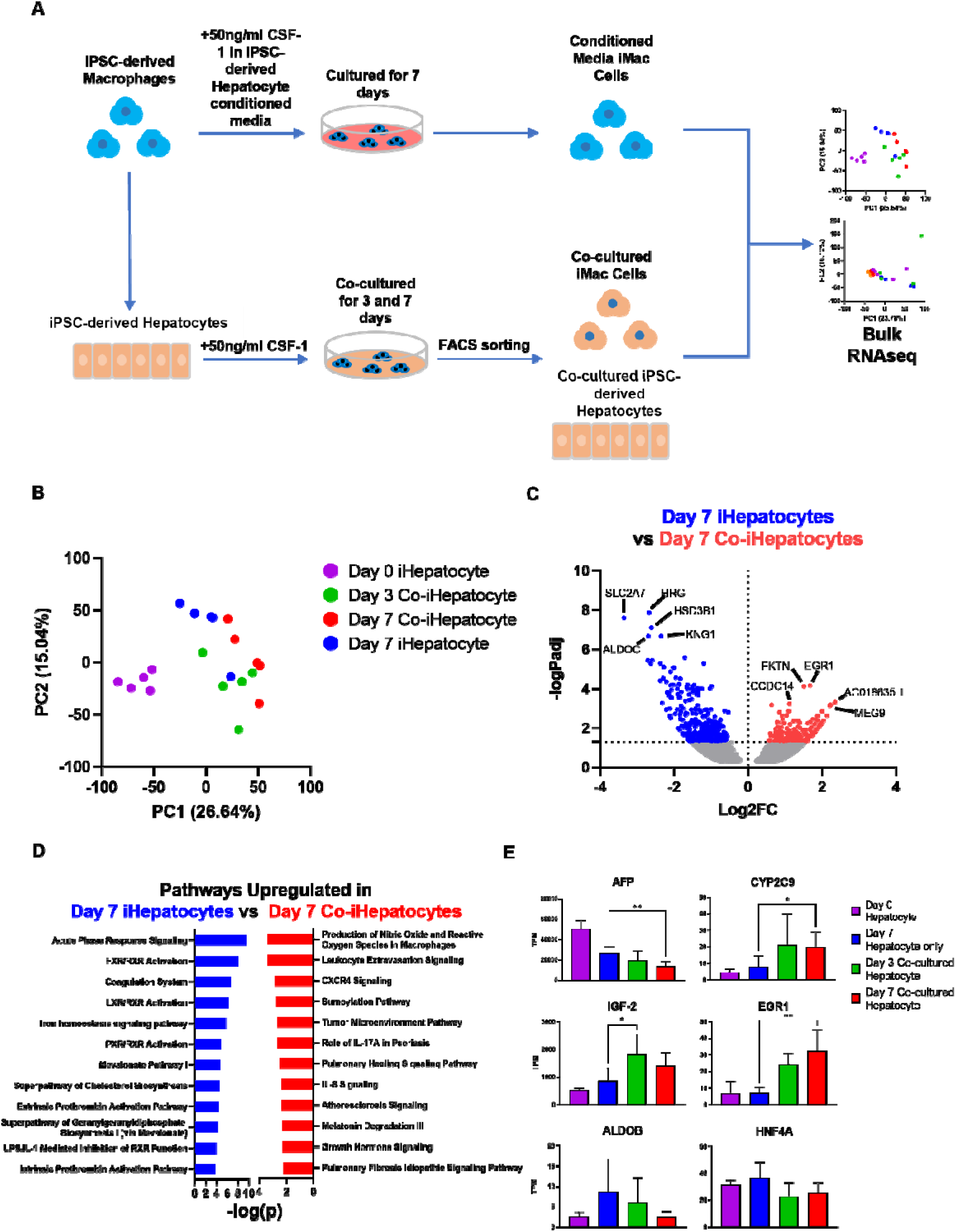
Model development and effects of iMacs on iHeps. A) Schematic of experimental layout. B) Principal Component Analysis of bulk RNAseq data from Day 0 iHeps, Day 3 co-iHeps, Day 7 co-iHeps and Day 7 iHeps. C) Volcano plot showing differentially-expressed genes between Day 7 iHeps and Day 7 co-iHeps. D) Differentially-expressed upregulated pathways between Day 7 iHeps and Day 7 co-iHeps. E) Gene expression levels of *AFP*, *CYP2C9*, *IGF-2*, *EGR-1*, *ALDOB* and *HNF4A* in Day 0 iHeps, Day 3 co-iHeps, Day 7 co-iHeps and Day 7 iHeps. Student’s T test was applied. *p<0.05, **p<0.01

### 2.4 Differentiation of human peripheral blood monocytes (PBMCs) into monocyte-derived macrophages

Human PBMCs were purified from blood apheresis cones by ficoll gradient (Cytiva, 17-1440-02), before being magnetically sorted with CD14+ microbeads (Miltenyi, 130-050-021). 1 x 10^6 CD14+ cells were then seeded per well of a 6 well plate in RPMI 1640 (Hyclone, SH30027.01) with 10% FBS (Biowest, S1810-500), 1x Pen/Strep, 1x Sodium Pyruvate, 1x NEAA and 1x L-Glut with 50ng/ml CSF-1 and cultured for 7 days. The purity of the monocyte-derived macrophages was then determined by flow cytometry.

### 2.5 Co-culture of iMacs and monocyte-derived macrophages with iHeps

2 x 10^5 iMacs or monocyte-derived macrophages were added to 5 x 10^5 iHeps during a full media change (4ml) per well of a 6 well plate on Day 0. We chose this ratio as it has been previously described that in the adult mouse liver, Kupffer cells can be up to 35% of hepatocyte numbers [15]. This media was comprised of Williams’ Medium E containing the following supplements: Penicillin/Streptomycin (10,000 U/mL / (10,000 μg/mL) – final conc = 1%, ITS+ - final conc= 2%, 4mM glutamax, 30mM HEPES buffer. The concentrations of the supplements were double that of standard conditions used due to the infrequent media change (please see section 3.1 for more details). In addition, 50ng/ml of CSF-1 was added to each well. The cells were cultured for 7 days, with a half-media change (2ml) on Day 4 consisting of the same media as described above with 100ng/ml of CSF-1, resulting in a final concentration of 50ng/ml CSF-1 in the well. The cells were collected by digesting with Accutase (StemCell, 07920) for flow cytometry, RNAseq analysis and qPCR. For pharmacological testing, the number of iHeps and iMacs were downscaled to 96-well plates while maintaining the same iHep:iMac ratio (2.5:1).

### 2.6 Flow cytometry

Standard staining procedures were used to prepare the cells for flow cytometry analysis. Briefly, cells were dissociated into single cells using the different methods described above for the various tissues, before being incubated with 100ul of antibody containing FACS buffer (1% BSA and 4mM EDTA in PBS) per 5 million cells for 20 minutes at 4°C. The cell suspension was then washed with 5ml of FACS buffer, centrifuged at 400g, and the supernatant was removed. The cells were finally resuspended in PBS containing 3uM DAPI (Invitrogen, D1306). Data was acquired by LSRII (BD Bioscience) and analyzed by Flow Jo (Tree Star, Inc.). For cell sorting, cells were sorted using FACS Aria II (BD Bioscience) or FACS Aria III (BD Bioscience). The following antibodies were used: FITC-conjugated anti- human CD14 (Biolegend, 325604), APC/Cy7-conjugated anti-human CD45 (Biolegend, 368516), PE-conjugated anti-human CD163 (R&D Systems, FAB1607P-100), Alexa Fluor 647-conjugated anti-human CX3CR1 (Biolegend, 341608, PE/Cy7-conjugated anti-human CD11b (eBioscience, 25-0118-42).

### 2.7 Bulk RNA-seq

Total RNA was extracted using Arcturus PicoPure RNA Isolation kit (Thermofisher, KIT0204) according to manufacturer’s protocol. All RNAs were analyzed on Labchip GX system (Perkin Elmer, United States) for quality assessment with median RNA Quality Score of 9.15. cDNA libraries were prepared using 2ng of total RNA and 1ul of a 1:50,000 dilution of Ambion ERCC RNA Spike-in Controls (Thermofisher, 4456740) using the SMARTSeq v2 protocol described [16] with the following modifications: 1. Use of 20mM TSO, 2. Use of 250pg of cDNA with 1/5 reaction of Nextera XT kit (Illumina, FC-131). The length distribution of the cDNA libraries was monitored using DNA High Sensitivity Reagent Kit (Perkin Elmer, CLS60672) on the Labchip GX system (Perkin Elmer). All samples were subjected to an indexed PE sequencing run of 2×51 cycles on HiSeq 2000 (Illumina) at 16 samples per lane.

### 2.8 Bulk RNA-seq analysis

Paired-end raw reads were mapped to GRCh38 human genome build using STAR aligner [17]. Genes were counted for reads that were mapped to genes using featureCounts [18] based on GENCODEv29 gene annotation [19]. Log2 transformed counts per million mapped read (log2CPM) and log2 transformed reads per kilobase per million mapped reads (log2RPKM) were computed using edgeR Bioconductor package [20]. Across all samples, genes with log2CPM inter-quartile range (IQR) less than 0.5 were filtered out from subsequent differential expression gene (DEG) analysis. DEG analyses for culture condition comparisons were all done using edgeR. Selection of DEGs was done with Benjamini-Hochberg [21] which adjusted P-values <0.05. R function ‘prcomp’ was used to perform Principal Component Analysis (PCA) on log2RPKM values. The pseudobulk values were calculated using the ‘AverageExpression’ function in Seurat. The pearson correlation were calculated using the ‘cor’ function in R. The GO enrichment was calculated using the ‘enrichGO‘ function from the ClusterProfiler package.

### 2.9 Quantitative real time PCR (qPCR)

Total RNA extracted using RNeasy Plus Micro-kit (Qiagen, 74034) was quantified using a NanoDropTM ND-1000 Spectrophotometer and converted to cDNA using iScript cDNA synthesis kit (Bio-Rad Laboratories, 1708890). qPCR was performed in 7000 Fast Real-Time PCR System (Applied Biosystems, Foster City, USA) with FastStart Universal SYBR Green Master (Rox) (Roche, 04 913 850 001) and primers from GeneCopoeia, Inc. (Rockville, MD, USA). Gyceraldehyde-3-phosphate dehydrogenase (GAPDH) served as internal control. Accession numbers of tested genes are listed in Table below:

| Target Gene | Gene Name | Accession |
| --- | --- | --- |
| GAPDH | glyceraldehyde-3-phosphate dehydrogenase | NM_001256799.2 |
| AFP | Alpha fetoprotein | NM_000295 |
| AAT | Alpha-1 antitrypsin | NM_001134 |
| ASGPR | Asialoglycoprotein Receptor | NM_001671 |
| ALB | albumin | NM_000477.3 |
| CYP1A2 | Cytochrome P450 1A2 | NM_000761.5 |
| CYP3A4 | Cytochrome P450 3A4 | NM_017460.5 |
| CYP2B6 | Cytochrome P450 2B6 | NM_000767.4 |
| CYP2C9 | Cytochrome P450 2C9 | NM_000771.3 |
| UGT1A3 | UDP-glucuroosyltransferase 1A3 | NM_019093.2 |
| GST1A2 | Glutathione S-Transferase | NM_000846.3 |
| MRP2 | Multidrug resistance-associated protein 2 | NM_000392 |
| CD14 | Cluster of differentiation 14 | NM_001174105.1 |
| CD32 | Cluster of differentiation 32 | NM_001136219.1 |
| CD68 | Cluster of differentiation 68 | NM_001251.2 |
| CD163 | Cluster of differentiation 163 | NM_203416.3 |
| ID1 | Inhibitor of DNA-binding protein 1 | NM_002165.3 |
| ID3 | Inhibitor of DNA-binding protein 3 | NM_002167.4 |

### 2.10 Enzyme-linked immunosorbent assay (ELISA) for measurement of cytokines

Interleukin-6 (IL-6) levels in the media were measured human IL-6 ELISA kit (Abcam, ab178013) according to manufacturer’s instructions. The cytokine production from dosed cells was normalized with the viability of cells measured from the same well. This ensures that changes in cytokines are not due to changes in cell numbers that might arise upon drug treatment. The normalized cytokine level was expressed as percentage of DMSO+LPS which allowed better comparison between different batches of experiments. This normalization approach has been used in previous studies [22].

### 2.11 Cell viability assay

Cell viability was examined with AlamarBlue^TM^ cell viability assay (Thermo Fisher Scientific Inc., Dal1025) according to manufacturer’s instructions. Briefly, AlamarBlue was diluted 10-fold using PBS containing 2 mg/mL of glucose and this working solution was added to the cells and incubated for 1 hour. Fluorescence (Ex: 530nm, Em: 590nm) readings were obtained using Tecan Microplate Reader M1000 PRO.

### 2.12 Drug administration

Following set up of co-culture as described in 2.5, the model was subjected to treatments including 100 ng/ml LPS (Sigma Aldrich, L2654), and drugs (Sigma Aldrich) (with/without LPS) for 48hr. Stock solutions of drugs were prepared in dimethyl sulfoxide (DMSO) and diluted in medium prior to use. Medium containing 0.1% DMSO was used as vehicle control except. After 48 hr, supernatant was collected cytokine measurement and cell viability in the same wells were measured as described in 2.11.

### 2.13 Statistical analysis

Mean and standard deviation were obtained from at least three independent batches of cells. Unpaired, paired student’s T-test and one-way or two-way analysis of variance (ANOVA) were performed accordingly at an overall confidence level of 95% using Prism software (GraphPad, San Diego, CA, USA) and indicated below each figure.

### 2.14 Data Availability

Raw data from RNA-seq analysis have been deposited in the NCBI Gene Expression Omnibus under accession number GSE307755.

## 3. Results

### 3.1 Optimization of culture conditions allow survival and function of iHeps and iMacs

In order to investigate the effects of co-culturing iHeps with iMacs, we first established and optimized a co-culture system utilising IMR90 iPSC-derived iHeps and iMacs (**Figure 1A**). To ensure survival and functionality of both cell types in our desired culture period (at least one week), certain key factors had to be accounted for: Hepatocytes are high in metabolically activity and often require frequent media change for replenishment of nutrients and removal of waste. On the other hand, it would be ideal to allow the iHeps and iMacs to interact through direct crosstalk as well as through secreted soluble factors without removing these factors from the system via media exchange. To assess the performance of hepatocytes with more infrequent media changes, expression of key hepatic markers was assessed when media was changed every other day, every two days or every three days and compared to a daily media change (**Supplementary Figure 1**). To compensate for the lower frequency of media change, we tested the effects of 2X supplements; standard media volume (**Supplementary Figure 1A**) and 2X supplements with 2X standard culture volume (**Supplementary Figure 1B**). Gene expression was compared as fold change to standard hepatocyte culture conditions (daily media change with 1X supplement and standard media volume). Increasing supplements alone could maintain the expression of ALB (albumin), AAT (alpha-1-antitrypsin), cytochrome P450 (CYP)- 1A2, 3A4, 2B6 and UDP-glucuronosyltransferase (UGT)1A3, but not CYP2C19 (36 - 68% decrease), glutathione S-transferase (GST)A2 (35-66% decrease) and multidrug resistance protein-2 (MRP2; 7 to 18% decrease) (**Supplementary Figure 1A**). In contrast, combining the increase in supplements with increase in culture volume resulted in maintenance of all hepatic markers (**Supplementary Figure 1B**). Albumin and urea production were also maintained to 1-1.4 pg/cell/48 hrs and 126-146 pg/cell/48 hrs (**Supplementary Figure 1C**) which is in the range of albumin and urea production reported by us [11, 14, 22] and others [23–27] previously.

### 3.2 Co-culture with iMacs improves iHep maturation and development

Cells sorted via flow cytometry on Day 3 and Day 7 (**Supplementary Figure 1D, 1E**), and the co-cultured hepatocytes (co-iHeps) and co-cultured iMacs (co-iMacs) were then analysed by RNA sequencing in to assess if co-culturing leads to transcriptomic changes reflecting improve differentiation and maturation. We also analysed Day 0 and Day 7 mono-culture controls, as well as iMacs cultured in PHCM for 7 days as control groups.

Principle component analysis (PCA) of the iHep samples revealed that that although the direction of change was the same for both the mono-cultured and co-cultured iHeps, there was a greater degree of change along the PC1 axis for co-iHeps (**Figure 1B**). We hypothesised that this might be due to the iMacs promoting further maturation and development of the iHeps, and performed a Pearson correlation between the mono-cultured and co-cultured iHeps against *in vivo* fetal liver hepatocytes from a published dataset [28], which revealed that the co-cultured iHeps were better correlated with the *in vivo* hepatocytes than the mono-cultured iHeps (**Supplementary Figure 1F**). Indeed, looking at the differentially expressed genes (DEGs) between the Day 7 iHeps and Day 7 co-iHeps revealed that the co-iHep upregulated genes associated with tissue morphogenesis and repair like Early Growth Response 1 (*EGR1*) [29], centrosome duplication such as the long non-coding RNA *CCDC14* [30] and angiogenesis-related gene like Maternally Expressed Gene 9 (*MEG9*) [31] (**Figure 1C, Supplementary Table 1**). On the other hand, iHeps cultured alone upregulated genes that inhibit proliferation and angiogenesis such as Histone Rich Glycoprotein (*HRG*) [32] and Kinogen 1 (*KNG1*) [33] (**Figure 1C, Supplementary Table 1**). We then performed pathway analysis using Ingenuity and discovered that the top upregulated pathway in Day 7 iHeps was acute phase response signalling, with the associated genes mainly relating to protein production such as Transthyretin (*TTR*) and Serpin Family D Member 1 (*SERPIND1*) (**Supplementary Table 2**), while the top upregulated pathway in Day 7 co-iHeps was related to nitric oxide and reaction oxygen species production, consisting mainly of metabolic and phosphatase genes like *c-FOS* and Serine/Threonine protein phosphatase 2A regulatory subunit B” beta (*PPP2R3B*) (**Figure 1D, Supplementary Table 3**). A closer look at the pathways upregulated in Day 7 iHeps and Day 7 co-iHeps also revealed that the Day 7 co-iHeps had a stronger tissue development and anabolic signature, upregulating pathways associated with tumor microenvironment, fibrosis and growth hormone signalling, while the Day 7 iHeps upregulated iron and cholesterol pathways.

Finally, we compared the expression of key genes across all the iHep samples (**Figure 1E**). In line with our hypothesis that co-culturing iMacs with iHeps would improve the maturation of the iHeps, we found that alpha-fetoprotein (*AFP*), a marker that is upregulated during fetal development and then downregulated as hepatocytes mature [34] followed a similar trend in our culture system, with lower expression in the Day 7 iHeps as compared to the Day 0 iHeps. Furthermore, the expression levels of *AFP* in Day 3 co-iHeps and Day 7 co-iHeps was even lower than that of Day 7 iHeps. Likewise Hepatocyte Nuclear Factor 4 (*HNFA4*), a transcription factor upregulated in hepatic progenitor cells [35], trended toward reduced expression in the co-cultured iHeps, albeit non-significantly. Insulin-like growth factor-2 (*IGF-2*), a key growth factor in fetal development [36], was significantly upregulated in the co-cultured iHeps 3 days after co-culture, with expression diminishing at the 7 day mark. Cytochrome P450 Family 2 Subunit C member 9 (*CYP2C9*), a cytochrome involved in drug metabolism was also significantly upregulated in the co-iHeps, while Aldolase Fructose-Biphosphate B (*ALDOB*), an enzyme whose upregulation is associated with unregulated cell proliferation and poor cancer prognosis [37] had a non-significant trend towards downregulation. Taken together, our data suggest that co-culturing the iHeps with iMacs increased their maturation and promotes improved hepatocyte development.

### 3.3 Co-culture with iHeps imparts KC-like identity to iMacs

Next, we turned our attention towards the iMacs within our co-culture system to test if they acquired KC features. PCA analysis showed that the co-iMacs clustered much tighter together, while iMacs that were cultured alone or in hepatocyte conditioned media (cond-iMac) were much more scattered (**Figure 2A**). This suggests that co-culturing iMacs in direct contact with iHeps may help maintain the identity of the iMacs better, mirroring how *ex vivo* resident tissue macrophages (RTMs) rapidly lose their identity when removed from their tissue environment [38]. DEG analysis of all the iMac groups showed that both the Day 7 iMacs and the Day 7 cond-iMacs shared an activated and inflammatory profile, upregulating genes such as TNFSF18 [39] and ACP5 [40] (**Figure 2B, Supplementary Table 4**). Day7 iMacs had a more migratory profile, uniquely expressing Chemokine Receptor 7 (*CCR7*) and Chemokine Ligand 5 (*CCL5*), while Day 7 cond-iMacs uniquely expressed Dishevelled Associated Activator of Morphogenesis 2 (*DAAM2*), a developmental regulator of the WNT pathway [41].

**Figure 2.**
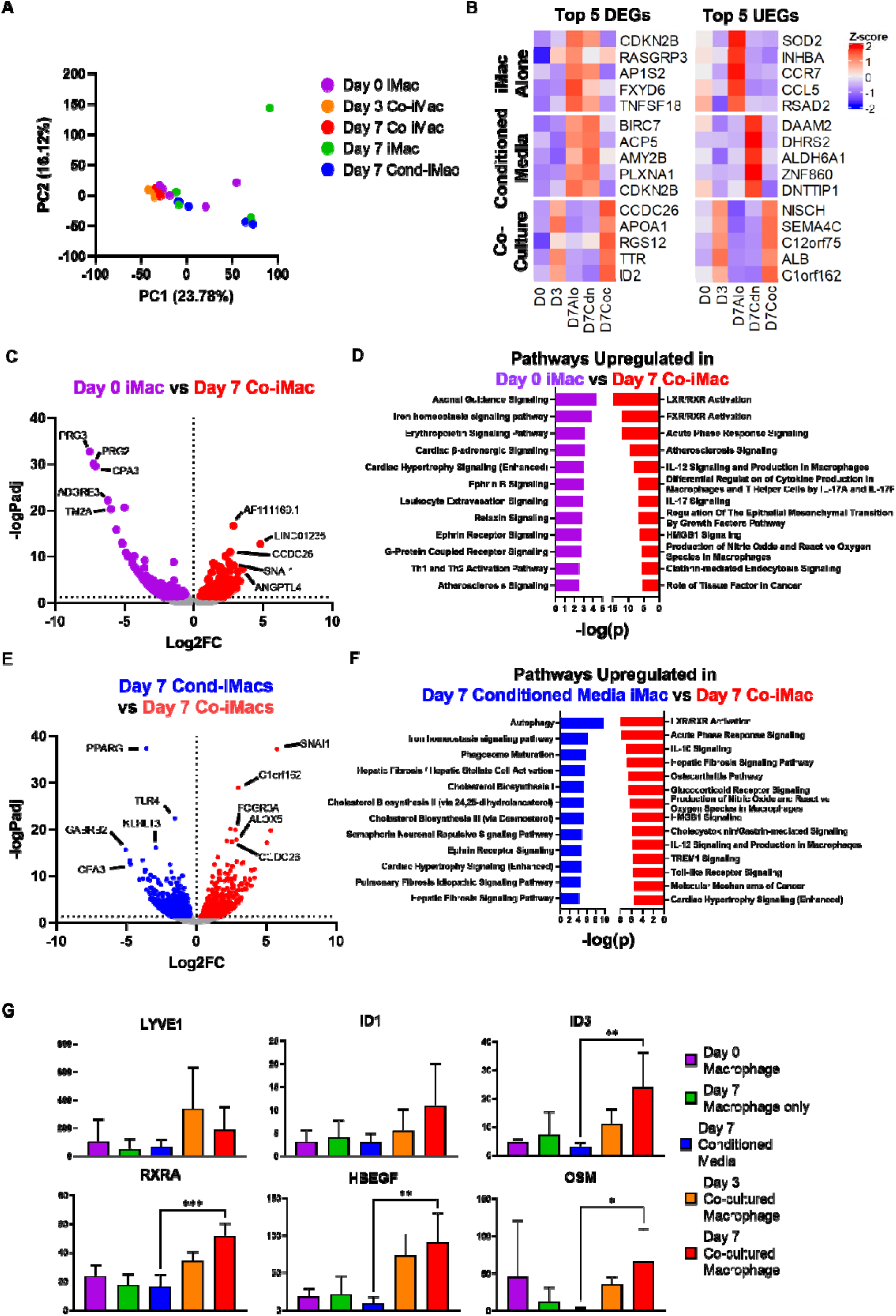
Effects of iHeps on iMacs. A) Principal Component Analysis of bulk RNAseq data from Day 0 iMacs, Day 3 co-iMacs, Day 7 co-iMacs, Day 7 iMacs and Day 7 cond-iMacs. B) Heatmap showing the top 5 differentially and uniquely expressed genes from culturing iMacs alone, with conditioned media (cond-iMacs) and after co-culture with iHepatocytes (Day 7 co-iMacs). C) Volcano Plot showing differentially-expressed genes between Day 0 iMacs and Day 7 co-iMacs. D) Differentially expressed upregulated pathways between Day 0 iMacs and Day 7 co-iMacs. E) Volcano Plot showing differentially-expressed genes between Day 7 cond-iMacs and Day 7 co-iMacs. F) Differentially expressed upregulated pathways between Day 7 cond-iMacs and Day 7 co-iMacs. G) Gene expression levels of *LYVE1*, *ID1*, *ID3*, *RXRA*, *HBEGF* and *OSM* in Day 0 iMacs, Day 3 co-iMacs, Day 7 co-iMacs, Day 7 iMacs and Day 7 cond-iMacs. Student’s T test was applied. *p<0.05, **p<0.01, ***p<0.001

As we were interested in how the presence of iHeps in such co-culture model might educate and impart KC-like identity to the iMacs, we looked at the DEGs between Day 0 iMac and Day 7 co-iMacs. Day 0 iMacs were more immunogenic, expressing Proteoglycan 2 Pro Eosinophil Major Basic Protein 2/3 (*PRG2/PRG3*) and Carboxypeptidase A3 (*CPA3*) (**Figure 2C**). In contrast, Day 7 co-iMacs upregulated the angiogenic Angiopoietin-like 4 (*ANGPTL4*) [42], matching the angiogenic signature observed in the Day 7 co-iHeps. Importantly, Ingenuity pathway analysis revealed that the top pathway in Day 7 co-iMacs is the Liver X Receptor/Farnesoid X Receptor (LXR/RXR) signalling pathway (**Figure 2D, Supplementary Table 5**), a key regulator of KC identity [43], while the top pathway in Day 0 iMacs is the axonal guidance pathway, consisting of chemokines like C-X-C Motif Chemokine Ligand 12 (*CXCL12*) and C-X-C Motif Chemokine Receptor 4 (*CXCR4*), metalloproteases like ADAM Metallopeptidase with Thrombospondin Type 1 Motif 15 (*ADAMTS15*), and semaphorins like Semaphorin 3C (*SEMA3C*) (**Supplementary Table 6**). This suggests that co-culturing iMacs with iHeps initiated a KC-specific programme, while the Day 0 iMacs had a more general and unspecified identity lacking environmental cues. In addition, Day 7 co-iMacs also upregulated pathways associated with tissue repair and remodelling, as well as tissue growth factors, while migratory and immune signalling related pathways were upregulated in the Day 0 iMacs (**Figure 2D**). Altogether, these data suggest that co-culturing iMacs with iHeps is sufficient to impart a more tissue supportive and KC-like identity to the iMacs.

The use of conditioned media as a surrogate of the *in vivo* tissue microenvironment is a popular approach, and we wondered how different iMacs cultured in direct co-culture with iHeps would be from iMacs cultured in conditioned media from primary hepatocytes (PHCM). Comparing the DEGs between Day 7 cond-iMacs and Day 7 co-iMacs, the Day 7 cond-iMacs upregulated immune-related genes such as Toll-Like Receptor 4 (*TLR4*) and *CPA3*, while the Day 7 co-iMacs more highly expressed Fc Gamma Receptor IIIa (*FCGR3A*), which is also upregulated in Kupffer Cells (**Figure 2E**). Pathway analysis showed that the top upregulated pathway in the Day 7 cond-iMacs was autophagy, which might indicate that conditioned media alone was insufficient to maintain proper macrophage biology in the absence of contact with other cells, although some hepatic related pathways were also upregulated as well (**Figure 2F, Supplementary Table 7**). On the other hand, the most highly upregulated pathway in the Day 7 co-iMacs was the FXR/RXR pathway, which along with the upregulation of other anabolic pathways related to hepatic health **(Supplementary Table 8)**, suggest that co-culturing rather than conditioned media is superior at inducing KC-like identity in macrophages.

We then looked at the expression of selected key KC genes. Lymphatic Vessel Endothelial Hyaluronan Receptor 1 (*LYVE1*), DNA-binding protein inhibitor ID1 (*ID1*) and DNA-binding protein inhibitor ID3 (*ID3*), which are important markers of KC identity [42]. *ID3* was significantly upregulated in the co-iMacs but not in the cond-iMacs, with a trend toward increased *ID1* and *LYVE1* expression as well (**Figure 2G**). Retinoid X Receptor Alpha (*RXRA*), the obligate binding partner of LXR [44], was also upregulated in only the co-iMacs. Interestingly, we also found that the co-iMacs upregulated the expression of Heparin Binding EGF Like Growth Factor (*HBEGF*) [45] and Oncostatin M (*OSM*) [46], which are potent effectors of liver regeneration and development.

Finally, we compared the iMacs from our study with *in vivo* embryonic human liver monocytes and KCs [47]. Pearson correlation revealed that regardless of condition the *in vitro* derived iMacs were poorly correlated to the *in vivo* human liver monocyte/macrophages (**Supplementary Figure 2A**), suggesting that either the co-culture duration was insufficient to induce full KC commitment or that there might be missing cues from other cell types that were not present in our co-culture. However, pathway analysis on the DEGs between the embryonic human liver macrophages and either the Day 7 cond-iMacs (**Supplementary Figure 2B, Supplementary Table 9**) or Day 7 co-iMacs (**Supplementary Figure 2C, Supplementary Table 10**) revealed that pathways relating to cytokine production, sensory development and eye development were differentially regulated between the Day 7 cond-iMacs and embryonic liver macrophages, while there were no significant pathway differences between the Day 7 co-iMacs and embryonic liver macrophages. This suggests that direct co-culture might drive the macrophages to adopt a more tissue developmental or supportive phenotype.

Altogether, the data supports our hypothesis that co-culturing iMacs with iHeps induces a KC-like tissue supportive identity in the iMacs, which in turn promotes the maturation and development of the iHeps.

### 3.4 Validation of the acquisition of KC-like identity by iMacs and maturation of iHeps in co-culture

We confirmed the observations from the RNA sequencing analysis by analysing gene expression of macrophage and KC-specific markers in Day 3 co-iMacs and Day 7 co-iMacs via RT-PCR. Macrophage markers *CD68*, *CD163*, *CD11b*, *CD32* and *CD14* were all upregulated at Day 3 and Day 7 by 1.8 to 7.6 fold compared to iMac mono-culture control (**Figure 3A**). Importantly, KC-specific markers *ID1* and *ID3* were upregulated by 3.8 fold and 3.9 fold respectively at Day 3 and by 6.3 fold and 6.5 fold respectively at Day 7 (**Figure 3B**). This suggests that macrophage phenotype is maintained during the 7 days of co-culture but KC-specific development increases in a time-dependent manner. When Day 7 co-iMacs were compared to iMacs cultured in PHCM (Day 7 cond-iMacs; without the presence of iHeps), macrophage marker upregulation compared to iMac mono-culture control were similar in both conditions (**Figure 3C**) but KC-specific *ID1* and *ID3* were upregulated to a lower extent in Day 7 cond-iMacs (2.6 fold and 2.5 fold respectively) compared to Day 7 co-iMacs (6.3 fold and 6.5 fold respectively; **Figure 3D**).

**Figure 3.**
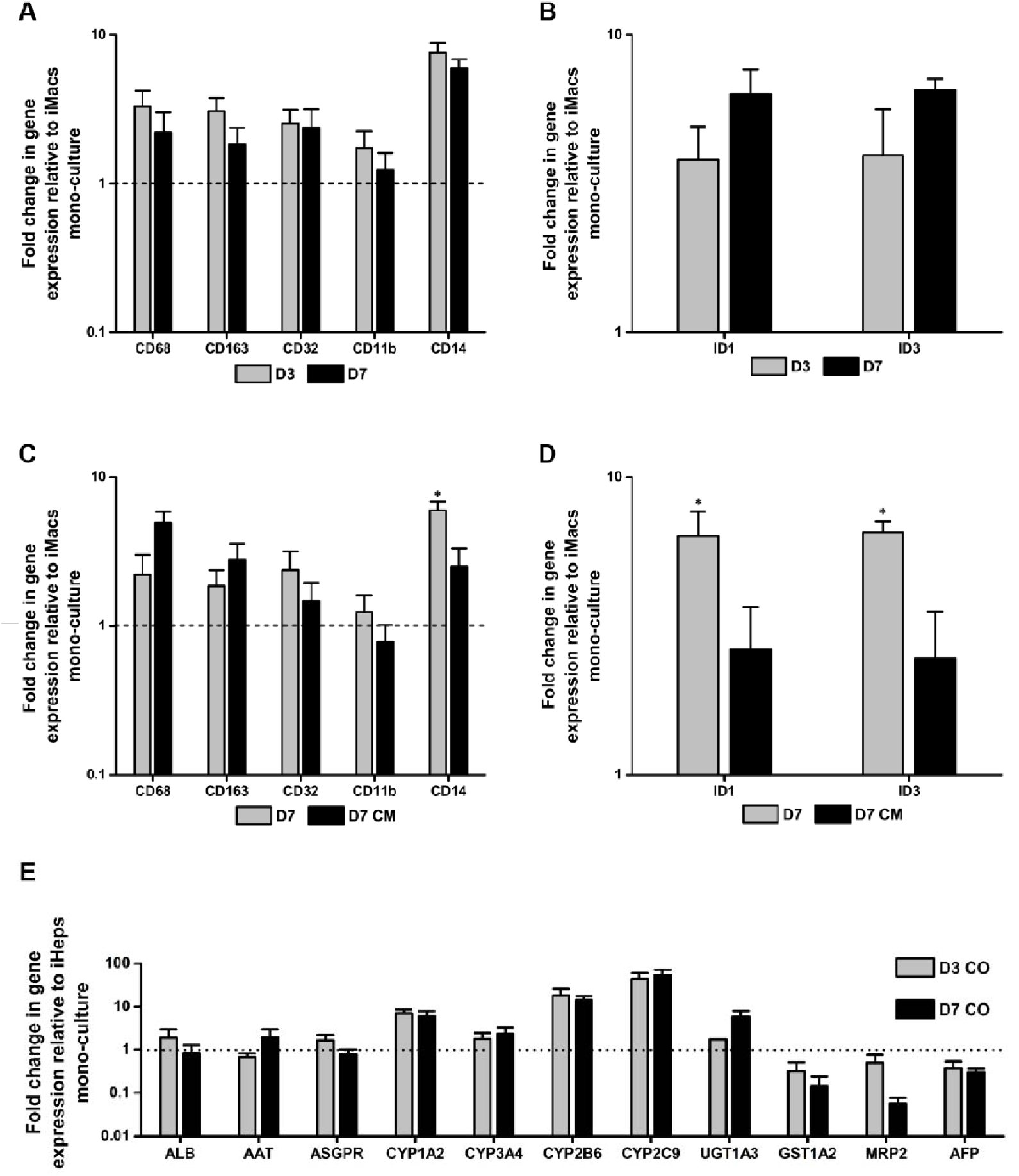
Gene expression of iMac and iHep markers in co-cultures. A) Expression of macrophage markers (left panel) and KC-specific markers (right panel) on day 3 and day 7 of co-culture. B) Expression of macrophage markers (left panel) and KC-specific markers (right panel) on day 7 of co-culture compared to iMac cultured in PHCM. C) Expression of hepatic markers (left panel) on day 3 and day 7 of co-culture. * p<0.05.

Next, we confirmed the RNA sequence analysis of iHep via RT-PCR. With the exception of Glutathione-S transferase1A2 (*GST1A2*) and Multidrug Resistance Associated Protein 2 (*MRP2*), co-culture with iMacs improved expression of several hepatic marker genes including *ALB*, α-1 antitrypsin (*AAT*), cytochrome P450 enzymes (*CYP1A2*, *CYP3A4*, *CYP2C9*, *CYP2B6*) and UDP-glucuronosyltransferases (*UGT1A3*) by 1.8-53.1 fold at Day 3 and Day 7 of co-culture (**Figure 3E**). Importantly, *AFP*, a marker that is expressed at high levels in fetal hepatocytes and increases with hepatocyte maturation [30], had 37% lower expression at Day 3 and 37% lower expression in Day 7 iHeps as compared to Day 0 iHeps (**Figure 3E**). This is consistent with RNA sequencing data that showed a downregulation of *AFP* (**Figure 1E**).

Overall, validation of the expression of key markers observed as modulated by RNA sequencing analysis confirms that culturing iMacs with iHeps imparts KC-like phenotype in iMacs and supports maturation and function of the iHeps.

### 3.5 Co-culture model accurately mimics clinical/*in vivo* immune responses of drugs

Drug-induced liver injury (DILI) is a complex and critical clinical problem with an estimated incidence rate of up to 19 cases per 100000 people in western countries [48]. In particular, immune-mediated idiosyncratic drug-induced liver injuries remains especially poorly understood, due in large part to the lack of appropriate human modelling systems [6]. To test the application of the co-culture in detecting immune-mediated response to hepatotoxic drugs, we carefully selected a group of 7 drugs based on the following criteria: 1) confirmed reports of immune/inflammation-associated effects according to 3 databases [49–51]; 2) available clinical data, or *in vivo* responses if the former is unavailable; and 3) known Cmax (maximum serum concentration of drug) to guide the concentration of drug used in the *in vitro* model. Drug concentrations used in our system was no more than 20-fold of its Cmax (Details of selected drugs are summarized in **Supplementary Table 11)**. The drugs included diclofenac (DIC), sulindac (SLD) leflunomide (LFM); amodiaquine (AQ), lamotrigine (LTG), penicillin (PEN), pyrazinamide (PZA). DIC [52], SLD [53] and LFM [54, 55] have been reported to cause a decrease in interleukin-6 (IL-6); whereas AQ and LTG have been reported to cause an increase in IL-6 in patients with drug-induced liver injury [51]. PEN and PZA were selected as negative controls as no immune-mediated effects of these drugs have been reported. The cultures were co-treated with lipopolysaccharide (LPS), which has been reported as an important initiating co-factor in the development of DILI [57].

Our co-culture model could correctly recapitulate IL-6 decrease in DIC (48%, 45% and 18% at 15.7 μM, 50 μM and 150 μM respectively), SLD (46%, 11% and 8% at 67 μM, 200 μM and 600 μM respectively) and LFM (65%, 46%, 38% at 32.5 μM, 125 μM and 500 μM respectively). IL-6 increase upon AQ (157-200 fold) and LTG (155-322 fold) treatment was also correctly recapitulated. No changes in IL-6 level in PEN and PZA-treated samples were observed (**Figure 4**). Importantly, when iMac-derived iKC-like cells were replaced with blood Monocytes/Macrophages, none of these cytokine changes were recapitulated with the exception of SLD treatment (**Figure 5**). This shows, from a functional/application perspective, how iMac-iKCs, but not MoMLJs, are indispensable for *in vitro* liver models that can detect immune-mediated changes of hepatotoxicants.

**Figure 4.**
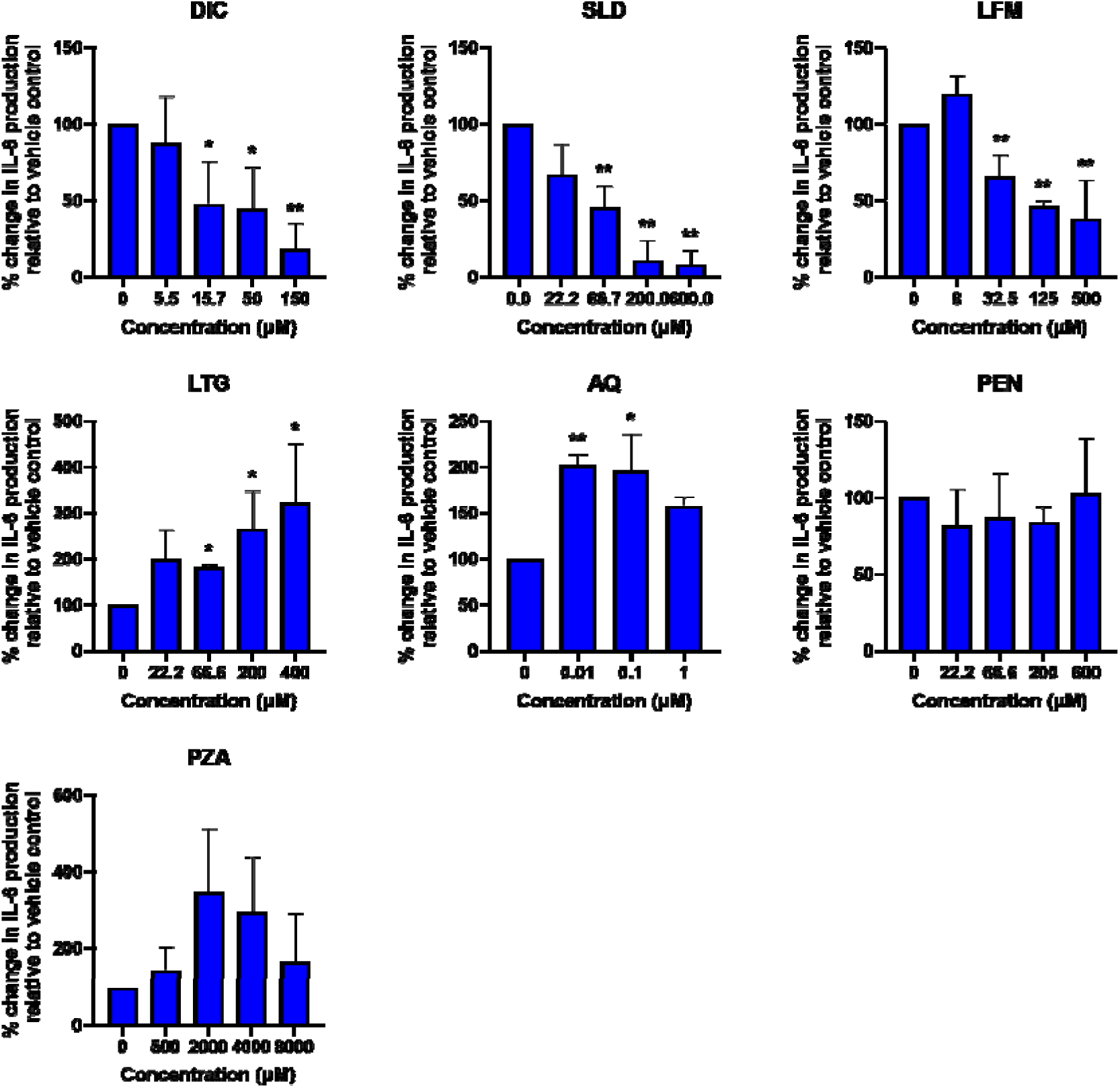
Changes in IL-6 level upon treatment with 7 paradigm compounds when iMac-derived KCs were used. Cytokine production was assessed in iMac-derived iKC and iHep co-culture treated with the drug stated in the plot title. Data is expressed as the percentage of the LPS-treated vehicle control. Error bars represented S.D., n=3. One-way ANOVA was applied. *p<0.05, **p<0.01 between treatment and vehicle control.

**Figure 5.**
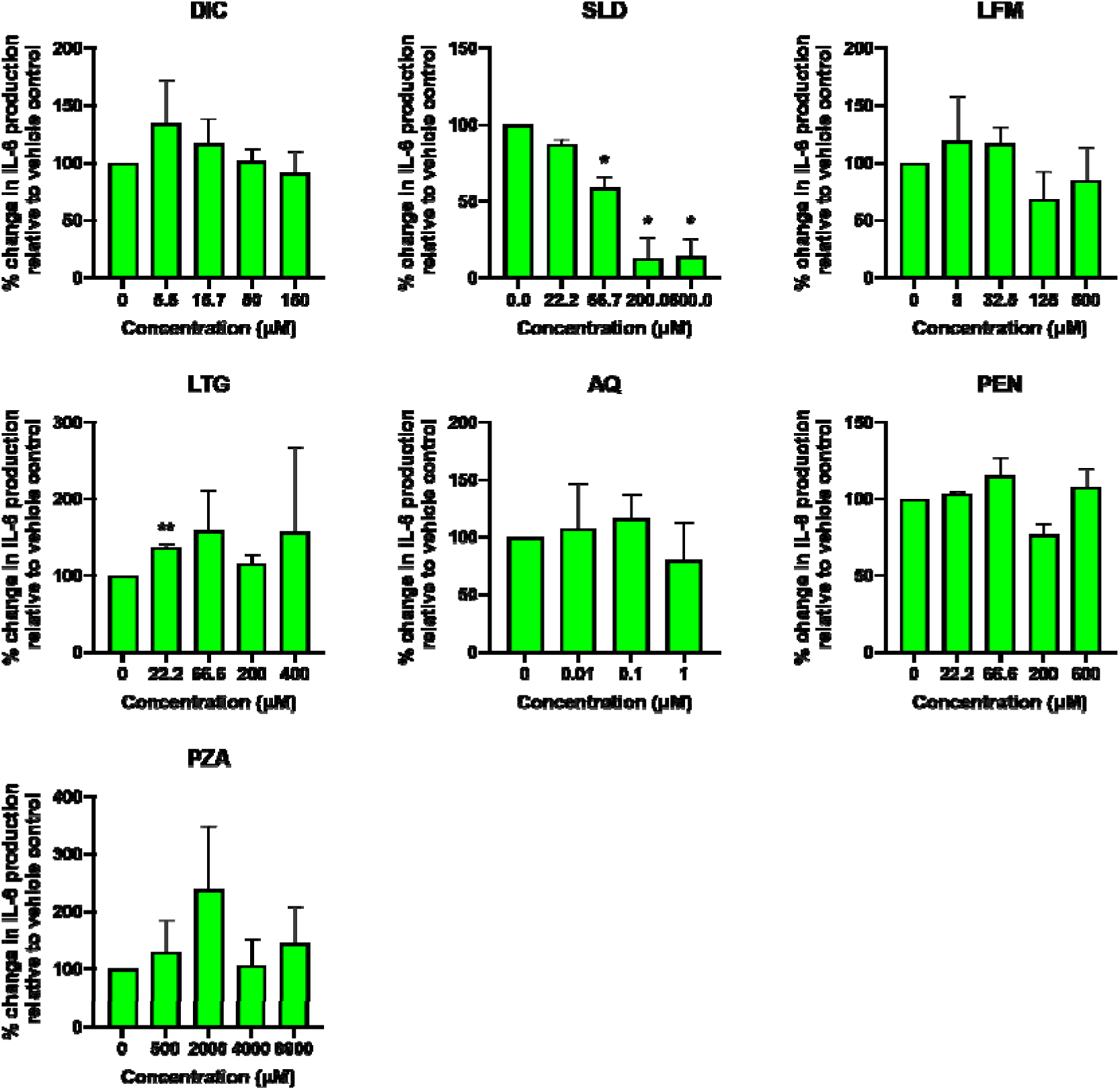
Changes in IL-6 level upon treatment with 7 paradigm compounds without iMac-derived KCs. Cytokine production was assessed in blood monocyte-derived macrophages and iHep co-culture treated with the drug stated in the plot title. Data is expressed as the percentage of the LPS-treated vehicle control. Error bars represented S.D., n=3. One-way ANOVA was applied. *p<0.05, **p<0.01 between treatment and vehicle control.

*In vivo*, KCs have been known to release mediators that are early features of inflammatory responses [56]. They are activated in response to inflammatory stimuli such as bacterial LPS that can damage the liver at large doses in a KC-dependent manner. At smaller doses, LPS activates KCs (without causing liver injury) but sensitizes the liver to a variety of other xenobiotics [57]. In our model, when the co-culture was co-treated with paradigm hepatotoxicants and LPS, drug-dose-dependent changes in cytokine production were observed (**Figure 4**), suggesting that the system can indeed mimic physiological responses, demonstrating the importance of liver-specific imprinting of macrophages in recapitulating responses in an *in vitro* liver model.

## 4. Discussion

IPSC-derived tissue cultures have tremendous potential in its ability to generate cells from all intra-embryonic lineages and have greatly expanded both our knowledge of mammalian tissue development and our technical toolbox for therapy and disease modelling [58]. However, a tissue is often more than the sum of its parts, and iPSC-derived tissue models have so far proven to be limited in their ability to effectively recapitulate the functionality and complexities of their presumed organs. In our study, we demonstrated how co-culturing iMacs with iHeps leads to not only acquisition of resident tissue characteristics within the iMac population, but also increased maturation of iHeps. We demonstrated that iPSC-derived Heps can directly impart hepatic macrophage-like identity to iMacs, giving them an iKC-like phenotype and genotype without the need for any additional soluble factors or components present in conditioned media. Application of this model was demonstrated by the ability of the system to mimic clinical IL-6 responses of 5 paradigm immune-mediated hepatotoxicants. Two additional hepatotoxicants with no known immune-mediated responses did not show any response in our model, demonstrating the specificity of the system. Finally, when the immune-mediated drug effects were tested in a system with iHeps and PBMC-derived macrophages, expected cytokine responses were not observed, indicating the importance of iKC-iHep co-culture in predicting drug mediated-immune responses.

Animal models have been used to model immune-mediated liver injury [59–61]; however, consistent liver injury was not observed upon treatment with drugs known to cause immune response in humans, even at high doses [62]. In cases where injury was observed, the characteristics were different from humans and possibly manifested through different mechanisms [61]. Such interspecies variations, contradictory findings as well as cost and ethical issues have spurred the development of *in vitro* models to supplement *in vivo* animal models. *In vitro* models, possibly due to the lack of appropriate human immune cell sources, have focused on soluble factor-driven single hepatocyte cultures in combination with inflammatory mediators or animal cell co-cultures [6]. However, mono-cultures lack cell-cell interactions, and animal *in vitro* models often fail to recapitulate *in vivo* and clinical findings. *In vitro* hepatocyte-KC models have previously been used; however most of them have utilised animal cells [63,64] which often prove inadequate in recapitulating *in vivo* or clinical findings. In order to avoid interspecies variations and improve the predictivity of *in vitro* models, appropriate human models still need to be developed and characterized.

The main challenge in studying human KCs in *in vitro* models is the cell source. The source of commercial primary human KCs (PhKCs) is limited and costly and high donor-to-donor variability is often observed [65]. Alternative human cell sources include THP-1 cells and peripheral monocyte-derived macrophages; however their cytokine profiles are dramatically different from human KCs [66, 67] and they lack liver-specific imprinting [43]. Even when human KCs were used, the work mostly focused on co-culture dependent improvement of hepatocyte function [6, 68], rather than interaction between the two cell-types and immune-mediated effect of drugs. Moreover, these models are often non-isogenic, which further confounds the inflammatory readouts. Hence, in our study, we aimed to first determine if the presence of hepatocytes could impart a tissue-resident-like phenotype to iMacs and drive them to be more Kupffer cell-like, then measured the systemic response to drug treatment compared to using monocyte-derived macrophages. While all tested drugs showed expected changes in cytokine production in the iMac-iHep co-culture (**Figure 4**), these changes were not recapitulated when the iMacs were replaced by MoMLJs (**Figure 5**) with the exception of SLD. Interestingly, expected IL-6 increase upon treatment with LFM could not be recapitulated in our previous model of iHep-iKCs co-culture where iKCs were generated using conditioned media from primary hepatocytes (data not shown). However, in our current model, LFM treatment resulted in the expected increase in IL-6, suggesting the potential importance of direct cellular cross-talk in iKCs differentiation and function.

Development of a suitable model system is further complicated by the understanding that KCs are derived from embryonic macrophages that seed the liver early on during development [7–9]. This also implies that conventionally used human peripheral blood MoMLJs might not be able to fully mimic KC biology. Thankfully, iMacs more closely resemble embryonic macrophages than MoMLJs [10] and as demonstrated in our own experiments, were able to acquire features of KCs upon co-culture with iHeps. Soluble factors were only partially responsible for this tissue adaptation, as in contrast to the iMacs that were cultured in hepatocyte conditioned media, the co-cultured iMacs upregulated genes in the LXR/RXR pathway, which has been shown to be important for KC specification and identity [43], further emphasising the importance of cell-cell contact for the acquisition of RTM identity. Furthermore, we also observed that these co-cultured iMacs were upregulating pathways associated with tissue repair and development, which might explain the increased maturity of the iHeps within the co-culture system. This is particularly relevant not only for hepatic model systems, but for other iPSC-derived model systems in general, as one of the key limitations of these systems is the functional and phenotypic immaturity of the differentiated cells [69]. Co-culturing iMacs with these immature iPSC-derived tissue model systems might provide the missing developmental cues needed for more physiologically and functionally relevant patient and disease models as well.

However, the results presented in our study are not without their own limitations as well. We primarily utilised bulk sequencing data in our analysis, and although we have demonstrated that there is minimal contamination between the hepatocyte and macrophage compartments of our co-culture, we are unable to completely rule out the minor possibility. Furthermore, our co-culture model is limited to only hepatocytes and macrophages, and might not fully address the complexity of the liver microenvironment. For instance, our model does not include other important cell types such as hepatic stellate cells [78] or sinusoidal endothelium cells [79]. This could perhaps explain why the day 3 co-cultured iHeps showed greater similarity to their i*n vivo* counterparts than the day 7 co-cultured iHeps, as the two-cell type co-culture alone might be enough to initiate co-development but insufficient to maintain it. Indeed, we observed transient upregulation of *IGF2* in the day 3 co-cultured iHeps (**Figure 1E**), suggesting the activation of a fetal developmental program. We are still unable to determine if its transient nature is an inherent feature of liver development or due to insufficient signalling. Finally, we also only specifically looked at *IL-6* expression as a systemic readout of liver function within our co-culture system. Although we were able to recapitulate the differential drug-specific expression of *IL-6* observed in various *in vivo* animal models of DILI [49–55], we did not explore the mechanisms behind the individual drug responses. It is important to note that a simplistic two-cell co-culture model system is still insufficient to fully encompass the complex systemic, metabolic and genetic landscape of a patient-specific condition such as DILI, but should instead serve to highlight the importance of incorporating other secondary cell types into model systems for a more holistic and accurate representation of tissue activity. Nevertheless, we believe that the development of co-culture systems containing multiple distinct cell types are an exciting next step in furthering our understanding of proper tissue specification and maturation, not only as a model of development but also tissue function.

---

**Supplementary Figure 1.**
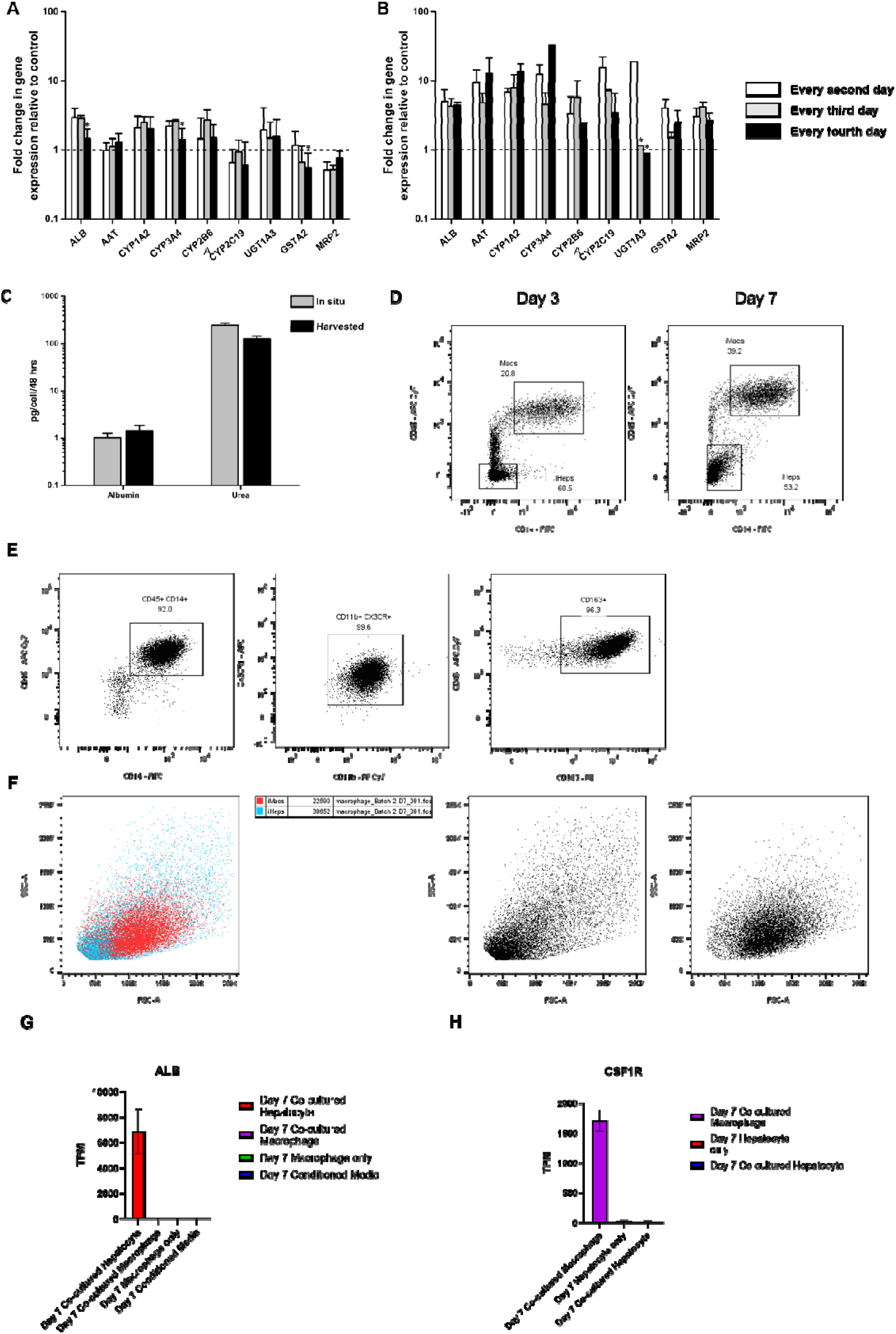
Optimisation of co-culture conditions and validation of purity. A & B) Fold change in expression of key hepatocyte genes in normal media volume and supplement concentration (A) or double media volume and supplement concentration (B), normalised to normal hepatocyte culture condition. C) Production of Albumin and Urea in co-culture system after 2 days, normalised to non-co-cultured hepatocytes. D) Flow cytometry of co-culture 3 and 7 days after co-culture. E) Flow cytometry showing purity of macrophages by CD45, CD14, CD11b, CX3CR1 and CD163 before co-culture. F)Flow cytometry showing relative cellular size and granularity of iMac and iHep populations after co-culture. G) Expression of Albumin in Day 7 co-cultured iHeps vs iMacs. H) Expression of CSF1R in Day 7 co-cultured iMacs vs iHeps.

**Supplementary Figure 2.**
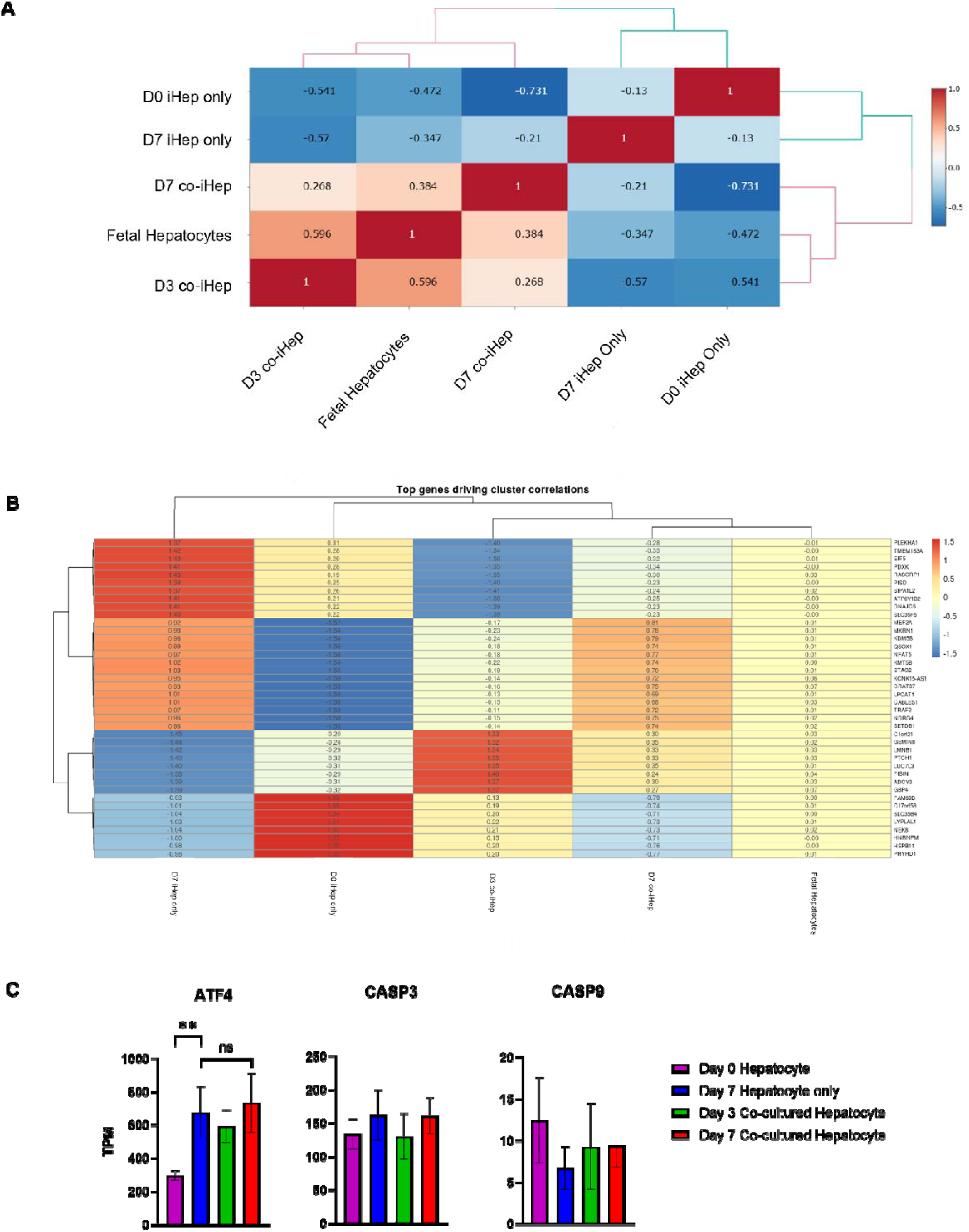
Correlation of iPSC-derived hepatocytes with or without co-culture with iPSC-derived macrophages. A) Pearson correlation of iHeps against fetal hepatocytes from Popescu et al, (2019). B) Top 40 genes influencing Pearson correlation of Supp Fig 2A. C) Expression of ATF4, CASP3 and CASP9 showed no significant upregulation of stress or apoptosis genes after co-culture.

**Supplementary Figure 3.**
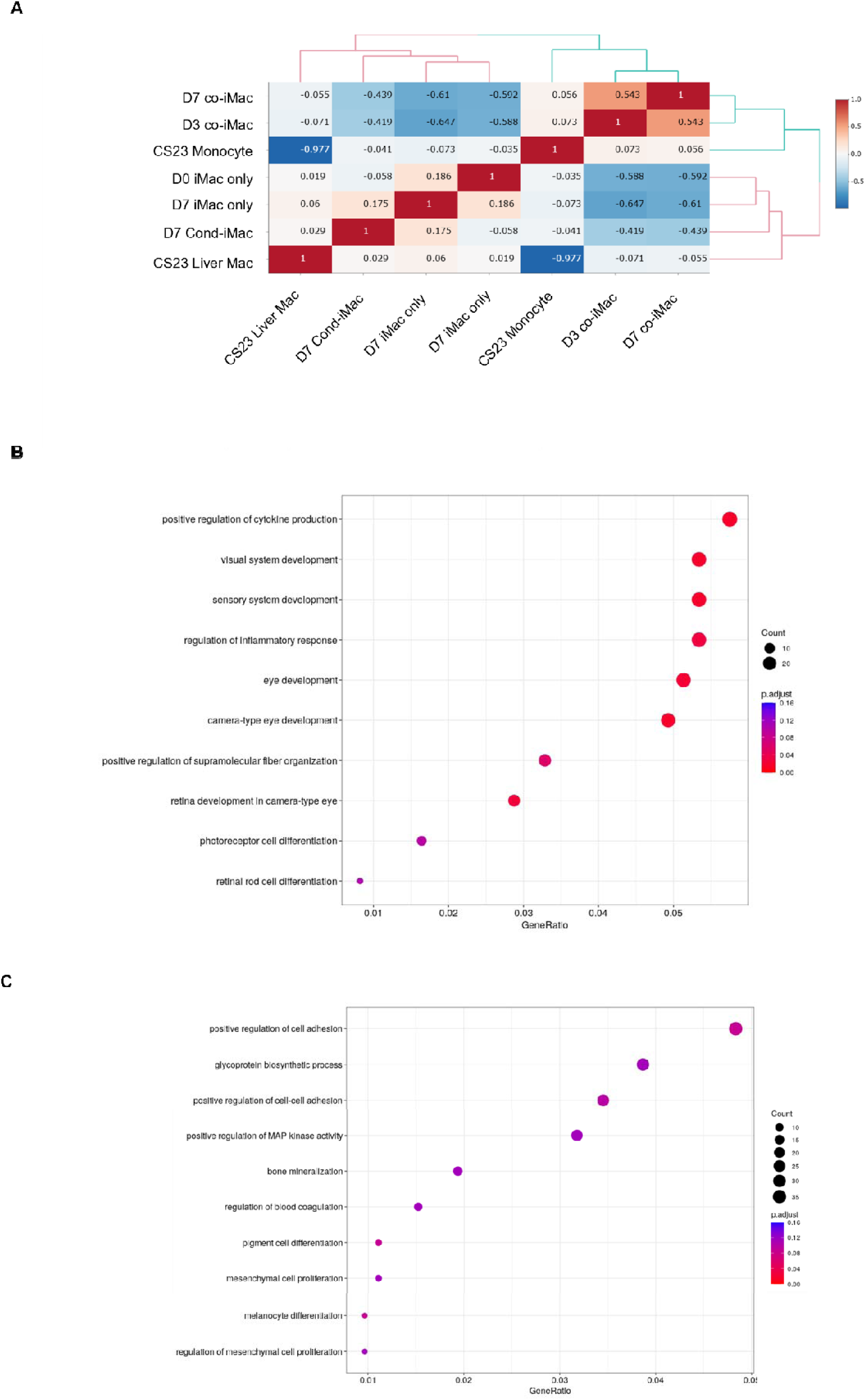
Correlation of iMacs with embryonic liver monocytes and macrophages, and pathway analysis of the DEGs. A) Pearson correlation of iMacs under control, conditioned media or co-culture conditions against embryonic liver monocytes and macrophages from Bian et al, (2020). B) Pathway analysis of DEGs between Day 7 conditioned media iMacs and embryonic liver macrophages. C) Pathway analysis of DEGs between Day 7 co-cultured iMacs and embryonic liver macrophages.

## List of Supplementary Tables

**Supplementary Table 1:** Differentially Expressed Genes between the iHeps Samples

**Supplementary Table 2:** Ingenuity Pathway Analysis for genes upregulated in Day 7 iHeps vs Day 7 co-iHeps

**Supplementary Table 3:** Ingenuity Pathway Analysis for genes upregulated in Day 7 co-iHeps vs Day 7 iHeps

**Supplementary Table 4:** Differentially Expressed Genes between the iMac Samples

**Supplementary Table 5:** Ingenuity Pathway Analysis for genes upregulated in Day 7 co-iMacs vs Day 0 iMacs

**Supplementary Table 6:** Ingenuity Pathway Analysis for genes upregulated in Day 0 iMacs vs Day 7 co-iMacs

**Supplementary Table 7:** Ingenuity Pathway Analysis for genes upregulated in Day 7 cond-iMacs vs Day 7 co-iMacs

**Supplementary Table 8:** Ingenuity Pathway Analysis for genes upregulated in Day 7 co-iMacs vs Day 7 cond-iMacs

**Supplementary Table 9:** GO enrichment results for DEGs between Day 7 conditioned media iMacs and embryonic liver macrophages

**Supplementary Table 10:** GO enrichment results for DEGs between Day 7 co-cultured iMacs and embryonic liver macrophages

**Supplementary Table 11:** Details of drugs used for testing

