## Supplementary material for "Direct contact between iPSC-derived macrophages and hepatocytes drives reciprocal acquisition of Kupffer cell identity and hepatocyte maturation": Supp Table 11

| Drug | Cmax/ (μM) | DILIrank [64] | DILIst [65] | Known immune-mediated effects [43-45] | Reported human responses | Reported in vivo (murine) responses |
|---|---|---|---|---|---|---|
| DIC | 8.0 [70] | most | 1 | ✓ | IL-6↓ [52]<br>IL-10↑ [52] | |
| SLD | 32.0 [70] | most | 1 | ✓ | Mechanistic IL-6↓ [53] | TNFα↑ [73] |
| LFM | 24.8 [74] | most | 1 | ✓ | | Cytokines↓ via NF-kB [54] |
| LTG | 9.8 [75] | most | 1 | ✓ | IL-6↑[56] | |
| AQ | 0.1 [76] | / | 1 | ✓ | IL-6↑ [56] | |
| PEN | / | less | 1 | × | | |
| PZA | 406.9 [77] | less | 1 | × | | |
